# Broad geographical and ecological diversity from similar genomic toolkits in the ascomycete genus *Tetracladium*

**DOI:** 10.1101/2020.04.06.027920

**Authors:** Jennifer L. Anderson, Ludmila Marvanová

## Abstract

The ascomycete genus *Tetracladium* is best known for containing aquatic hyphomycetes, which are important decomposers in stream food webs. However, some species of *Tetracladium* are thought to be multifunctional and are also endobionts in plants. Suprisingly, *Tetracladium* sequences are increasingly being reported from metagenomics and metabarcoding studies of both plants and soils world-wide. It is not clear how these sequences are related to the described species and little is known about the non-aquatic biology of these fungi. Here, the genomes of 24 *Tetracladium* strains, including all described species, were sequenced and used to resolve relationships among taxa and to improve our understanding of ecological and genomic diversity in this group. All genome-sequenced *Tetracladium* fungi form a monophyletic group. Conspecific strains of *T. furcatum* from both aquatic saprotrophic and endobiont lifestyles and a putative cold-adapted clade are identified. Analysis of ITS sequences from water, soil, and plants from around the world reveals that multifunctionality may be widespread through the genus. Further, frequent reports of these fungi from extreme environments suggest they may have important but unknown roles in those ecosystems. Patterns of predicted carbohydrate active enzymes (CAZyme) and secondary metabolites in the *Tetracladium* genomes are more similar to each other than to other ascomycetes, regardless of ecology, suggesting a strong role for phylogeny shaping genome content in the genus. *Tetracladium* genomes are enriched for pectate lyase domains (including PL3-2), GH71 *α*-1,3-glucanase domains and CBM24 *α*-1,3-glucan/mutan binding modules, and both GH32 and CBM38, inulinase and inulin binding modules. These results indicate that these fungi are well-suited to digesting pectate and pectin in leaves when living as aquatic hyphomycetes, and inulin when living as root endobionts. Enrichment for *α*-1,3-glucanase domains may be associated with interactions with biofilm forming microorganisms in root and submerged leaf environments.

## INTRODUCTION

The freshwater fungi traditionally called “aquatic hyphomycetes” are best known as ecologically important decomposers in stream ecosystems. The group is polyphyletic, its members are classified in various orders and families of Ascomycetes and Basidiomycetes. These fungi thrive on leaves and other plant debris that enter streams and other water bodies from terrestrial plants. In the process, carbon and nutrients from recalcitrant plant compounds, including cellulose and lignin, become accessible to diverse consumers in the stream food web (reviewed in (1, 2)). Up to 99% of the carbon in some streams enters as plant debris that is largely inaccessible to aquatic consumers without the degradative activity of these fungi (3, 4). Aquatic hyphomycete adaptations to stream environments include the production of asexual propagules (conidia) while under water. These spores are often branched, typically tetraradiate, or sigmoid, readily detachable from the fungus (*e*.*g*. from conidiophores) and passively dispersed in water. The conidia strongly adhere to surfaces upon contact. These features facilitate dispersal and colonization of new substrates (5-8). Less well known is that some aquatic hyphomycetes are dual niche (9) or multifunctional (10) species that differ in mode of nutrition or lifestyle in different environments, and are both decomposers of dead plant material in water (saprotrophs) and endobionts in terrestrial or aquatic plants (6, 9, 11-15). Although it has been known since at least 1939 (16) that some aquatic hyphomycetes could be isolated from terrestrial sources, these fungi are still intensively studied in their aquatic context due to their massive presence ecological importance, especially in streams.

The genus *Tetracladium* de Wild. (Ascomycetes, Leotiomycetes) was described in 1893 as “*parmi les algues, les diatomées et les débris de végétaux supérieurs dans les étangs, les fossés*” [among algae, diatoms and higher plant debris in ponds, ditches] (17). Since that time, conidia of *Tetracladium* have been recorded in aquatc habitats world-wide (18). The genus has grown to include eleven species, most of which produce distinctive, characteristically shaped conidia (19) (Fig. 1). The three most recent additions to the genus, *T. ellipsoideum, T. globosum*, and *T. psychrophilum* are the exceptions (20). These species were described from glacial soil from the Tibet Plateau and have elliptical and globose spores in the first two cases, and no reported spores in the third. These are the first species in the genus described from soil. However, it has been known since 1970 that *Tetracladium* could be isolated from terrestrial sources including sterilized roots (*T. marchalianum* (21)) and one species, *T. nainitalense*, was described from endophytic isolates (11). *Tetracladium*-like fungi, fungi identified as *Tetracladium* based on ITS similarity without morphological information, are increasingly reported to be among the most frequently found and abundant taxa in culture and metagenomics based studies of soil fungi, rhizosphere fungi, and endophytes world-wide. These studies represent diverse ecosystems, soils, and associated plants, ranging from domesticated crops (carrot (22) oilseed rape (23), ginseng (24), lettuce (25) and wheat (26, 27) to wild orchids *(28)* and mosses (29), from agricultural fields (30) to glacial and subglacial soils (31, 32), from sea level (33) to high altitudes (above 2600 m.a.s.l (34)), and spanning the globe from the Arctic (35) to the Antarctic (29, 36). Three psychrophilic species have been described from glacial soils from the Tibet Plateau (20)—one of the most extreme environments on earth; fungal extremophiles are of ecological and industrial interest due to their roles in ecosystem functioning and the secondary metabolites they produce (33, 37). Although hundreds of ITS sequences for *Tetracladium*-like fungi are available in public databases, it is not yet known how the fungi from these diverse environments are related to each other whether they fit within the genus *Tetracladium* as understood morphologically and phylogenetically at present. The generic type species, *T. marchalianum*, was neotypified in 1989 (38), but lack of the neotype ITS sequence also hinders taxonomic efforts within the genus.

**Figure 1:**
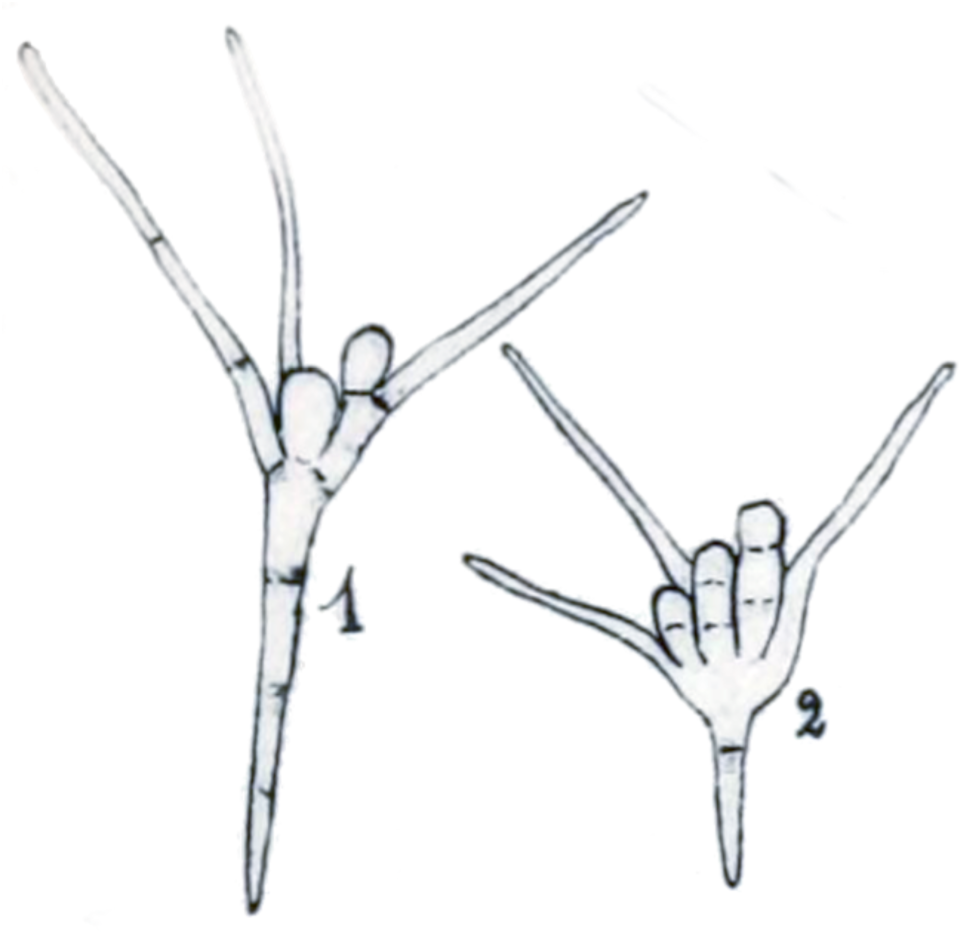
Conidia of *Tetracladium marchalianum* (1) and *T. cf. breve*(2), both originally described as *T. marchalianum*. Illustrations by de Wildeman (17)(b.1866-d.1947). Modified from plate IV(17). For reference, from the base of conidia 1 to the rounded terminus is around 35 µm.

Compared to the aquatic lifestyle of *Tetracladium* species (2), little is known about the biology or ecological importance of *Tetracladium* living outside of water. *In vitro*, some species are known to degrade lignin (39), pectin (40), startch (41), solubilize phosphate (42), have antimicrobial activity (43, 44), and to increase metal tolerance in host plants (45). Correlation studies have associated *Tetracladium* with increased plant growth (46), however, clear patterns are not yet emerging from which the roles or impact of *Tetracladium* as endophytes or in soil can be inferred (47). From studies in other fungal functional groups, such as pathogens and non-aquatic hyphomycete endophytes, there is increasing data available about secondary metabolite (SM, *e*.*g*. (48)) and carbohydrate active enzyme (CAZymes, *e*.*g*. (49)) profiles, as predicted from genomes, that may be associated with different lifestyles. By comparing the predicted molecular repertoires for *Tetracladium* both among strains from different ecological niches within the genus and to those of better studied functional groups it may be possible to gain insight into the ecology of *Tetracladium*. This reverse ecology approach to understanding *Tetracladium* biology has not before been possible; to date there are no publicly available *Tetracladium* genomes. Relatively few molecular tools exist for these species, they include microsatellites (50), mass spectrometry profiles (51), and taxon-specific fluorescence in situ hybridization probes (52).

The i) ecological importance of *Tetracladium* species in streams; ii) global reports of *Tetracladium*-like species in soils and as endophytes; iii) paucity of information about the non-aquatic roles of these fungi; iv) emerging reports of *Tetracladium* species from extreme environments; all signal a need to better understand fungi callasified in the genus *Tetracladium*. Moreover, these features raise an exciting suite of questions and opportunities for study. How have *Tetracladium* species evolved to tolerate environmental extremes? What are the features of genomes and their regulation that enable multifunctionality and broad environmental tolerance or adaptation in these species? Can the same genotypes or species thrive as endophytes and aquatics? How is multifunctionality distributed across the *Tetracladium* phylogeny? It is not possible to address all these questions in a single study, but here the foundations are laid to address them and to support advances across all aspects of *Tetracladium* biology. In this study, genomes of all known species of *Tetracladium* and several *Tetracladium*-like species were sequenced, with more than one representative strain per species when available. Phylogenomics was used to resolve evolutionary relationships within the genus. To gain insight into the biology of these fungi from their genomes, secondary metabolites and carbohydrate-active enzymes were predicted from the genomes to compare among species and with profiles for plant pathogens and other functional groups. Further, analysis of the ITS sequences of *Tetracladium*-like fungi from studies world-wide were analyzed together with the strains sequenced herein to test and illustrate the geographic and ecological diversity of the genus.

## MATERIALS AND METHODS

### Strains and sequencing

Twenty-four fungal strains, including all described species in the genus *Tetracladium*, were obtained for genome sequencing (see Table S1 for strain information and Nagoya Protocol status). *Tetracladium marchalianum* JA005 (full designation JA005.1) and *T. setigerum* JA001 are new collections from the Fyris River, Sweden. The remaining strains were obtained from the Czech Collection of Microorganisms (CCM), CABI, China General Microbiological Culture Collection (CGMCC), the Yeast Culture Collection at the Universidad de Chile, from Dai Hirose at Nihon University, Japan, and from Magdalena Grudzinska-Sterno from the Swedish University of Agricultural Sciences.

Cultures were maintained on potato dextrose agar (Fluka, cat #70139) at room temperature with ambient light or at 4°C in the dark (CGMCC strains) with backup cultures frozen at −80°C in 18% glycerol. Strains were grown in flasks with 50-200 mL of malt extract peptone medium (17 g/L malt extract, 2.5 g/L bacto-peptone) at room temperature with shaking for up to one month to obtain sufficient mycelium for DNA extraction. Mechanical disruption via grinding in liquid nitrogen or bead beating (Mini-Beadbeater Biospec 25/s, for 3 s or less) was performed on fresh or freeze dried mycelium before DNA extraction using the Zymo Quick DNA Fungal/Bacterial Miniprep Kit (D6005) as directed.

Sequencing libraries were prepared at the SNP&SEQ Technology Platform in Uppsala, Sweden, from 1 μg DNA using the TruSeq PCRfree DNA sample preparation kit (FC-121-3001/3002, Illumina Inc.) targeting an insert size of 350 bp as directed. Libraries were then sequenced paired-end with a 150 bp read length on an Illumina HiSEQX Ten, using v2.5 sequencing chemistry.

### Genome assembly and annotation

Sequence reads were quality and adapter trimmed using Trimmomatic v0.36 (53). Bases with quality scores below 3 were removed from the start and end of reads. Sequence falling below a quality score of 20 in a sliding window of size 4 and adapter sequences were removed. Resulting sequences greater than 36 bp were retained for genome assembly. Assembly was performed in in Spades v3.11.1 (54) using the “careful” option to reduce mismatches and short indels. The resulting scaffolds were used for further analysis. Assembled genomes were evaluated in QUAST V4.5.4 (55) and BUSCO v3.0.2b (56) using the Pezizomycotina odb9 database to test for completeness. Average depth of coverage was estimated in QualiMap v2.2.1 (57) after read mapping in BWA v0.7.17 (58).

Genomes were annotated using an iterative MAKER v3.01.2-beta (59) annotation pipeline. In the initial run, protein and EST evidence from *Botrytis cinerea* B05.10 (ASM83294v1), *Glarea lozoyensis* (ATCC 20868), *Phialocephala subalpine* (PAC v1), and *Sclerotinia sclerotiorum* (ASM14694v1), accessed from Genbank (28-29 May 2018), were used as evidence to train the Hidden Markov Model. Species specific repeat libraries generated using RepeatModeler v1.0.8 (60), using default settings and the NCBI search engine, were used for repeat masking. Outputs from this run were used to support *ab initio* gene predictions in GenMark-ES v4.33 (61) using the fungus algorithm option. The GenMark outputs were then used in MAKER for gene annotation on scaffolds larger than 1000 bp.

Putative gene functions and protein domains for the annotated genes were identified using Blastp (62) against the UniProtKB/Swiss-Prot annotated non-redundant database (63) (downloaded 29 October 2018) to return no more than one hit per gene with an evalue threshold of 1×10^−6^ and InterProScan v5.30-69.0 (64) with mapping to Gene Ontology and member database signatures (without using the pre-calculated match lookup service). These data were used to update the annotations in MAKER.

### Single copy orthologs and phylogenomics

Single copy orthologs (SCO) present in all Tetracladium strains and 27 additional ascomycetes (Table S2) were selected from ortholog groups identified by OrthoMCL v2.0.9 (65) (with inflation index 2). Annotations for *Cadophora malorum* and *Articulospora tetracladia* are not available at NCBI and were generated for use herein as described above. Before alignment, SCO sequences were processed in PREQUAL v1.02 (66) using default settings to remove sequence stretches with no evidence of homology. The SCO were then aligned (MAFFT v7.407 G-INS-I) with a variable scoring matrix and αmax=0.0.8 (67) and high-entropy regions were trimmed (BMGE v1.12 (68) with the BLOSUM95 similarity matrix). The resulting dataset (all-SCO) and two subsets of the data were used for analyses. The two subsets contained SCOs from the all-SCO dataset that are also in the BUSCO fungi (fungi-SCO) or ascomycete (asco-SCO) odb9 databases. Phylogenetic analyses were performed with maximum likelihood (ML) in IQ-TREE v1.6.8 (69). For the all-SCO dataset, a guide tree inferred under the X+G+C60 model (X = best-fit model inferred by BIC in ModelFinder (70); for +C60 see (71)). The guide tree and best-fit model from that analysis were used to infer the final tree with the posterior mean site frequency (PMSF) model (71) which better accounts for heterogeneity of amino acid frequencies across sites. Branch support was evaluated using ultrafast bootstrap (UFBoot, (72)) and SH-like approximate likelihood ratio test (SH-aLRT), each with 1000 replicates. The fungi-SCO and asco-SCO datasets were both analyzed using the best-fit models identified by ModelFinder and support estimated using 1000 UFBoot and SH-aLRT replicates. The fungi-SCO dataset was additionally analyzed using the FreeRate model (X+R5). Lifestyle, ecology and other designations for the fungi in this study are based on the actual sample sequenced, this can differ from the generalized biology of the species (Tables S1, S2). Trees were visualized and annotated using iTOL (73) with *Tuber borchii* as the root.

### ITS phylogeny

ITS sequences including the term *Tetracladium* were retrieved from GenBank and aligned in Geneious 10.2.4 (http://www.geneious.com). Sequences containing data spanning the ITS region (ITS1, 5.8s, and ITS2) were trimmed using BMGE (DNAPAM1 matrix). Using a distance matrix produced in Geneious, sequences with identical bases and alignments were identified and removed from the dataset. Sequences with excessive uncalled bases (>30) or poor alignments were also removed. The strains sequenced herein may be doubly represented in the dataset if not identified as duplicates in this process. Information about each sequence was manually searched to determine the origin and lifestyle when possible. The ITS-region ML phylogeny was determined in IQTree with the best-fit model and branch support estimated using 1000 non-parametric bootstrap replicates and visualized with *Penicillium antarcticum* as the root.

### Carbohydrate active enzymes (CAZymes) and secondary metabolites

To identify genes involved in recognition, metabolism, and synthesis of complex carbohydrates, the proteins annotated in MAKER for each strain were compared against existing CAZyme-related databases using the dbCAN2 meta server (74) (accessed online at http://bcb.unl.edu/dbCAN2, September-October 2019) with HMMER, DIAMOND, and Hotpep and the default settings. Only CAZymes predicted by at least two tools were used in further analyses. To count the domains identified, each prediction identifier for multi-family/multi-domain genes were counted separately not together as a new combined category. Secondary metabolite clusters were identified in genomes using the antiSMASH 5.0 fungal version public web server (75) (accessed online at https://fungismash.secondarymetabolites.org, August-September 2019) including use of the *KnownClusterBlast* function. Cluster analysis and visualization were done in R (76) on scaled (*scale*) count data using *pheatmap*. Differences in numbers and types of predicted CAZymes and SM among the genus *Tetracladium*, other Leotiomycetes, and the remaining taxa were evaluated using ANOVA in R, with taxon-group as the main effect, and post hoc analyses with Tukey’s HSD (*p* ≤ 0.05) as implemented in *agricolae*.

## RESULTS

### Strains

Four strains from this study have been deposited in the CCM culture collection and will be publicly available after peer reviewed publication; JA005 (CCM 9013), JA001 (CCM 9012), AFCN889 (CCM 9011), and AFCN900 (CCM 9010). *Tetracladium sp*. 82A210 failed to revive from preserved and long term cultures at the time the cultures were deposited. Efforts to revive this strain are ongoing and it will be deposited in the CCM if successful.

### Genomes

Twenty-four *Tetracladium* and *Tetracladium*-like strains, including all described species, were sequenced and the resulting raw data deposited as ENA Study PRJEB36440 (available after peer review). Data for each genome, including contigs, trimming, assembly, and coverage statistics as well as sample and read number details are provided in Table S3. The genomes assembled into 266 – 6,765 scaffolds (median = 378). Scaffold N50 values ranged from 185,943 – 2,082,751 bp (median = 894,004 bp). Total assembled genome sizes ranged from 34.3 – 43.6 Gb. Genome-wide average GC content in the assemblies was 46.13%. These values exclude scaffolds smaller than 500 bp. The *Tetracladium sp*. T11Df assembly had the lowest depth of coverage (69X) and had more than 5-times more scaffolds than the next highest, T. *maxilliforme* CCM F-529 with 1032. Depth of coverage ranged from 69 – 178X (average = 128X). These high depths of coverage were not targeted, rather they are the by-product of small genome size and high sequencer output; the 24 barcoded genomes were pooled and sequenced in one lane on an Illumina HiSeqX Ten. All genomes are highly complete as assessed in BUSCO. Of the 3,156 genes included in the Pezizomycotina odb9 database, 20 - 34 were missing from the genome of any strain. The genomes had 98.3 – 98.7% complete BUSCOs, and 97.8 – 98.5% were complete and single copy.

### Annotations and Orthologs

The number of annotated gene models ranged from 7769 in *T. sp*. T11Df to 9799 in *T. sp*. 82A210 (mean 8768, median 8694; Table S3). To predict gene function, the annotated genes for each strain were compared to the UniProtKB/Swiss-Prot annotated non-redundant protein database. These analyses returned similarity-based information for an average of 6810 (median 6759) predicted genes per strain. An average of 8681 protein signatures per strain (median 8613) were identified by comparing the annotated gene models to the InterProScan database. The gff files containing all annotation data will be available after peer review. A total of 1820 SCO present in all 51 genomes (all-SCO) included in this study were identified from the output of OrthoMCL.

### Phylogeny

The all-SCO dataset contained 794,205 aligned amino acid sites, 434,869 were parsimony-informative (Table S4). Analysis using the PMSF model starting with the best fit model LG+F+I+G4 and a guide tree (Fig. S1) as input, resulted in a tree with all branches fully supported (100% UFBoot and SH-aLRT)(Fig 2). All known and putative *Tetracladium* strains in this analysis form a monophyletic clade with three distinct groups. Group A contains all *Tetracladium* species with typical Tetracladium conidia, including *T. sp*. F-10008 which also produces conidia typical for Tetracladium when submerged (19). *Tetracladium sp*. 82A210, conidia yet unknown, was isolated as an endophyte of wheat and is resolved within *T. furcatum*. Group B includes the moss endobionts from the Antarctic, *T. sp*. AFCN889 and *T. sp*. AFCN900, which appear to be conspecific and the three soil species from the Tibet Plateau, *T. ellipsoideum, T. psychrophilum*, and *T. globosum*. Group C is monotypic, containing only the Antarctic yeast *T. sp*. T11Df. Only two branches in the initial guide tree were not supported (UFBoot values <95% and SH-aLRT <80%), one within Group A, separating *T. furcatum* from *T. maxilliforme*, and one within the Leotiomycetes separating *Phialocephala scopiformis* and *Articulospora tetracladia* (Fig. S1).

**Figure 2:**
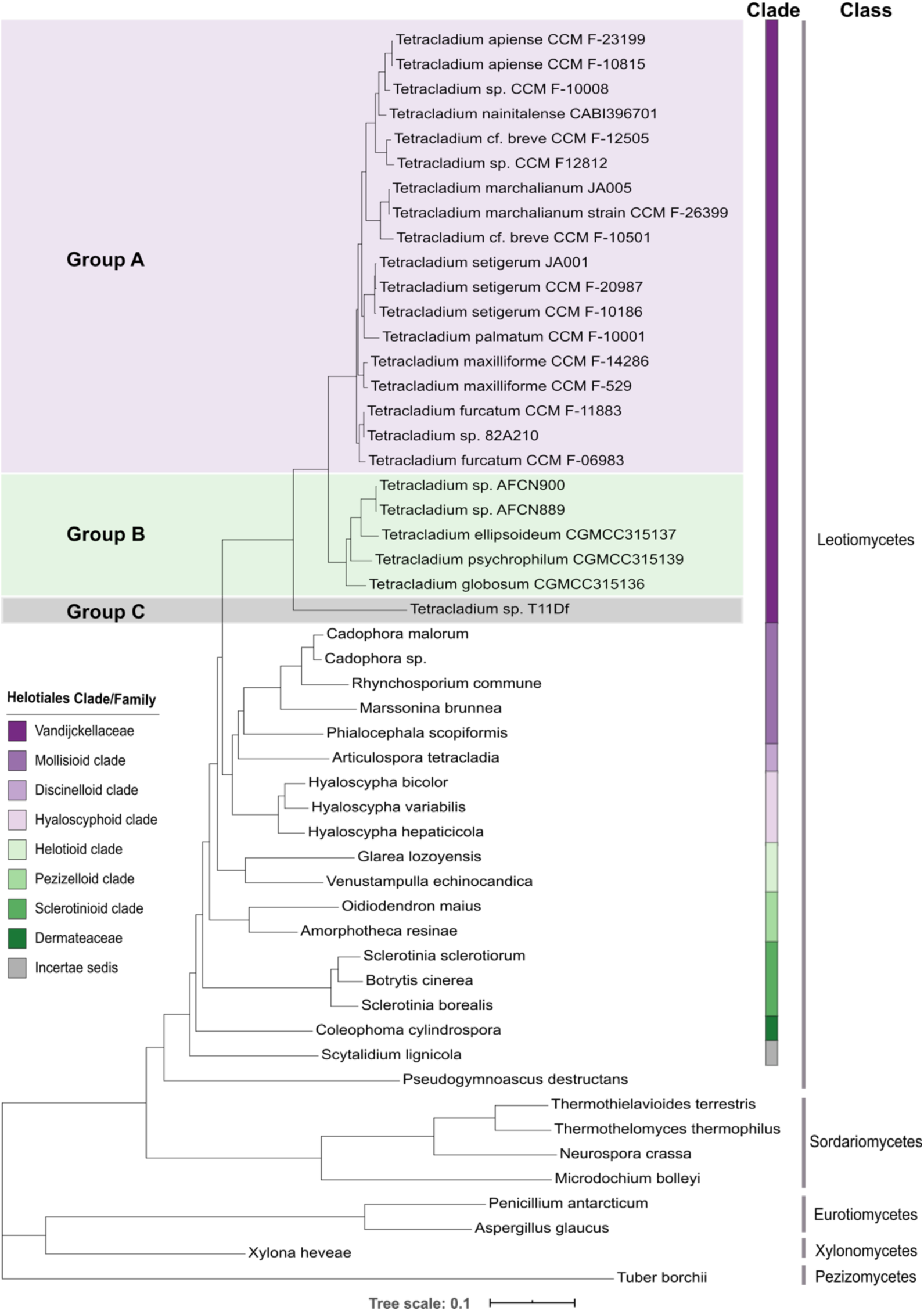
Phylogeny of *Tetracladium* and *Tetracladium*-like taxa (Groups A, B, and C) within the Leotiomycetes. Maximum likelihood phylogram inferred from 1820 single copy orthologs present in all 51 taxa in this analysis using the PMSF model in IQ-Tree. For fungi within the order Helotiales the color bar indicates the clade or family in which they belong, after Johnston *et al*. 2019 (details Table S2). All branches in this tree are fully supported (100% UFBoot and SH-aLRT). The tree scale represents the expected number of nucleotide substitutions per site. Phylogram drawn with *Tuber borchii* as the root.

To confirm the robustness of the phylogenetic relationships inferred from analyses of the large all-SCO dataset additional analyses were performed using subsets of the data with more complex substitution models than were computationally feasible for full dataset. The asco-SCO dataset contained 182,311 parsimony informative amino acid sites (339,460 total) from 709 SCOs (Table S4). ModelFinder identified LG+F+R6 as the best fit model. The resulting tree had a highly similar topology to the all-SCO result, with uncertain resolution for *T. furcatum/T. maxilliforme* and *P. scopiformis/A. tetracladia* (Fig. S2).

The fungi-SCO dataset was the smallest, with 35,673 parsimony informative sites (72,086 total) from 164 SCOs (Table S4). The best-fit model for the data was LG+F+R4. When analyzed using this model there was again poor resolution for *T. furcatum/T. maxilliforme* and *P. scopiformis/A. tetracladia* (Fig. S3). There was also insufficient support for the branch to *T. palmatum* and *T. setigerum*. Using the empirical mixture model (C60), all branches that were unsupported with the simpler LG+F+R4 model met the threshold for support, but the branch separating *Coleophoma cylindrospora* from *Scytalidium lignicola* was not supported (Fig. S3). Overall the phylogenies obtained in these analyses were highly consistent. Within the *Tetracladium* and *Tetracladium*-like clade, only the relationships of *T. furcatum* and *T. maxilliforme* to each other and the rest of Group A varied among analyses.

### ITS Phylogeny

To evaluate how ecological diversity is distributed within the genus, a phylogeny including 198 ITS DNA was produced and visualized in with information about the sequence (Fig. 3, Table S5). To further highlight diversity in the genus, orchid and challenging/extreme-environment associated sequences are also indicated. The later, identified as “cold” in Fig. 3, are from high latitude, high altitude, alpine, and glacier associated locales. The analysis included 229 sequences, with seven outgroups and the 24 strains sequenced herein. Of 402 total sites, 144 were parsimony informative and the best fit model for the data was TIM3e+R3. Branch support was determined using standard non-parametric bootstrapping (not UFBoot); bootstrap values >75% are considered well-supported using this approach. Conservatively, only branches with bootstrap values >85% are highlighted in the resulting figure (Fig. 3). As expected, this ITS tree does not reflect relationships between the species of *Tetracladium*, but similar ITS sequences from different sources and lifestyles are clustered. Only one *Tetracladium*-like sequence, JX630692, is not similar to species in Groups A, B, and C. When compared to other ascomycetes in GenBank, this sequence does not return hits to any culture identified *Tetracladium* species. Thus, the “*Tetracladium*” identity of this sequence is suspect.

**Figure 3:**
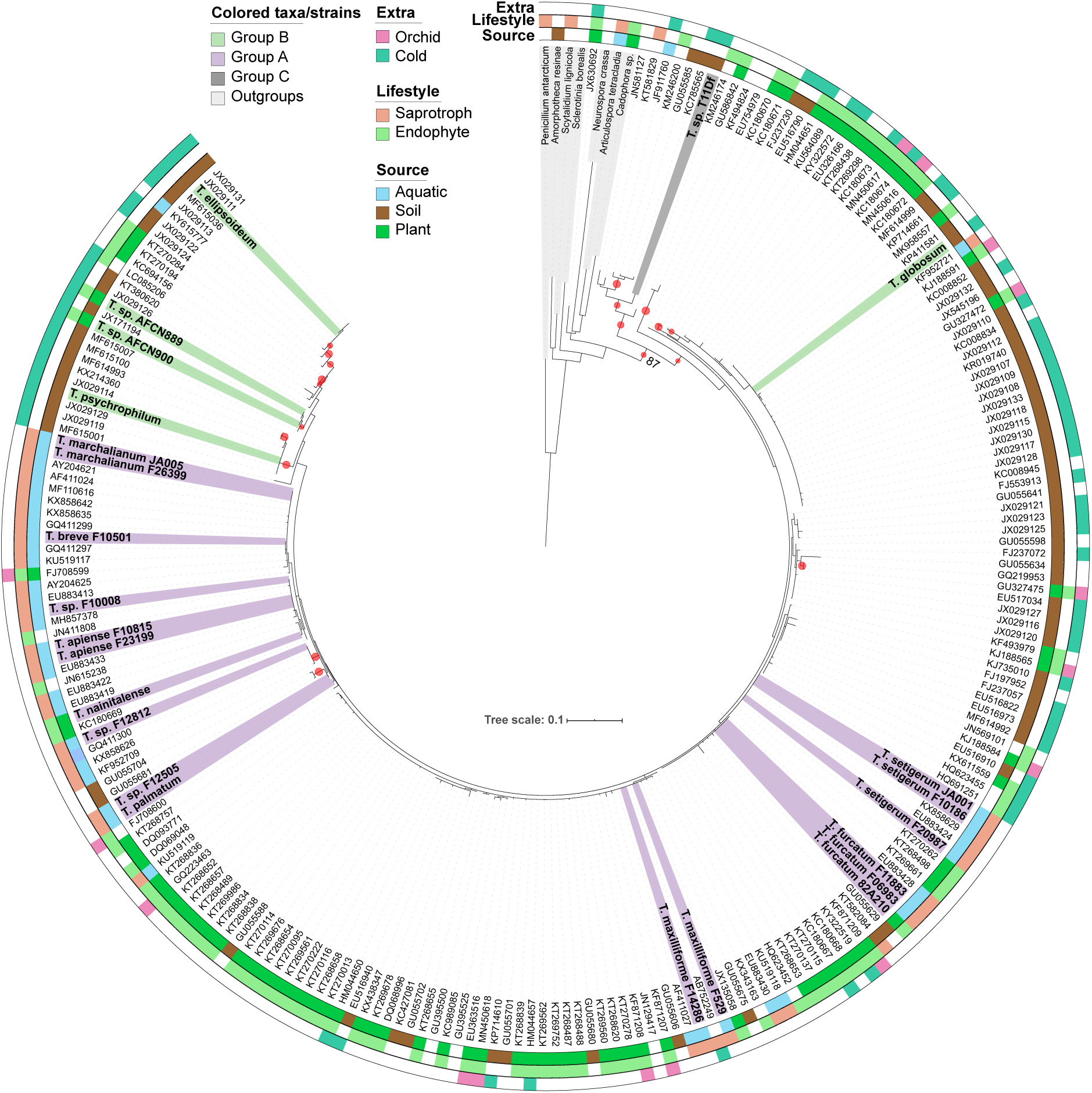
ITS and ecological diversity within the *Tetracladium* and *Tetracladium*-like fungi based on ITS region data and sample information for 198 accessions from GenBank, the 24 newly sequenced genomes herein (Groups A, B, and C colored as in Fig. 2), and outgroups (grey). Maximum likelihood phylogram inferred using the model TIM3e+R3. Note: Only branches receiving non-parametric bootstrap support > 85% are highlighted in this figure. Sequences identified as “cold” in the Extra category originate from high latitude, high altitude, alpine, or glacier associated locales. The tree scale represents the expected number of nucleotide substitutions per site. Phylogram drown with *Penicillium antarcticum* as the root.

### CAZymes

Cluster analysis based on the predicted number of carbohydrate-binding modules (CBM) and catalytic domains of each CAZyme class by species did not reveal any discernable ecological grouping (Fig. 4, Table S6). Clustering of the majority of the *Tetracladium* species reflects phylogeny, not ecology, because the only non-*Tetracladium* aquatic hyphomycete in the study does not cluster with *Tetracladium* species and the *Tetracladium* species cluster together regardless of ecology. The exception is the Antarctic yeast, *T. sp*. T11Df, which clusters with taxonomically and ecologically diverse species.

**Figure 4:**
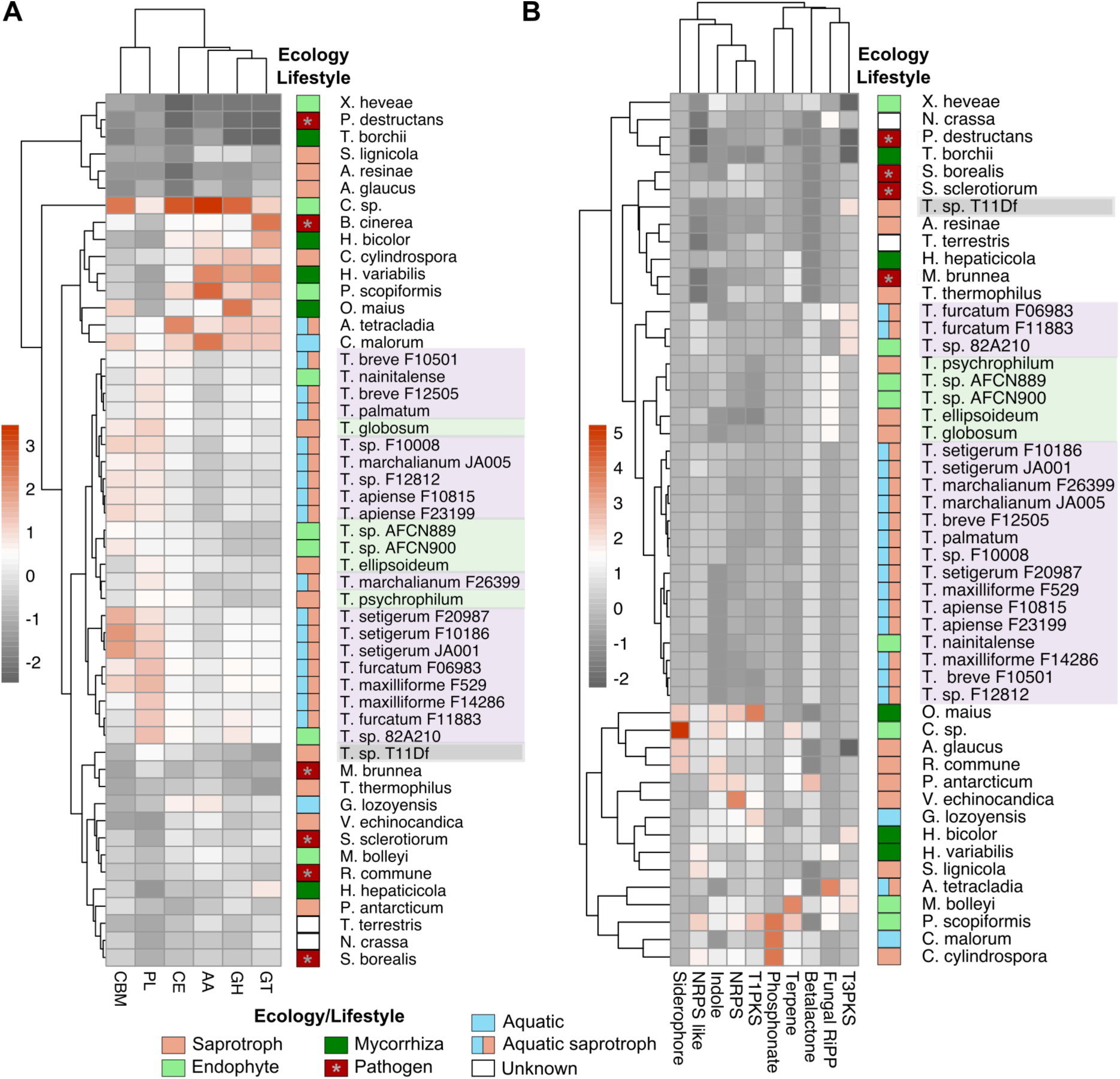
*Tetracladium* and *Tetracladium*-like strains cluster together in analyses of the number of CBM and CAZyme domains (A) and secondary metabolite clusters (B) of each type regardless of ecology. Heat map colors are relative to scaled datasets with orange colors representing high abundances and grey colors representing lower abundances of the domain/cluster in each genome. Note: *P. destructans* is an animal pathogen, all other pathogens infect plants. CAZyme classes and associated modules: Auxiliary Activities (AA), carbohydrate binding modules (CBM), carbohydrate esterase (CE), glycoside hydrolase (GH), glycosyltransferase (GT), and polysaccharide lyase (PL).

Like other Leotiomycetes (490.8 ± 161), *Tetracladium* genomes (487.6 ± 47.9) contain more CAZyme and CBM domains than other ascomycetes (329.1 ± 102.6; Fig. 5A; F_2_ = 6.97, *p* < 0.002; Tukey’s HSD, p ≤ 0.05). Likewise, Leotiomycetes and *Tetracladium* genomes have more predicted carbohydrate esterase (CE), glycoside hydrolase (GH), and glycosyltransferase (GT) domains than other ascomycetes (Fig. 5C; CE: F_2_ = 8.5, *p* <0.0001; GH: F_2_ = 5.9, *p* =0.005; GT: F_2_ = 6.3, *p* =0.004; Tukey’s HSD, p ≤ 0.05). However, *Tetracladium* genomes have fewer predicted Auxiliary Activities (AA) domains than other Leotiomycetes, and so are in line with other ascomycetes (F_2_ = 6.7, *p* = 0.0003; Tukey’s HSD, p ≤ 0.05). Note that the taxon-groups differ in sample size(*Tetracladium* = 24, Leotiomycetes = 18, Other ascomycetes = 8) and represent diversity at different taxonomic levels. Also, only domains predicted by HMMER are presented here, summaries of all HMMER results (Table S6) and results from Hotpep and DIAMOND (Table S7) are provided. These numbers are influenced by the contents of the databases and the genome assemblies and are thus “predicted” values.

**Figure 5:**
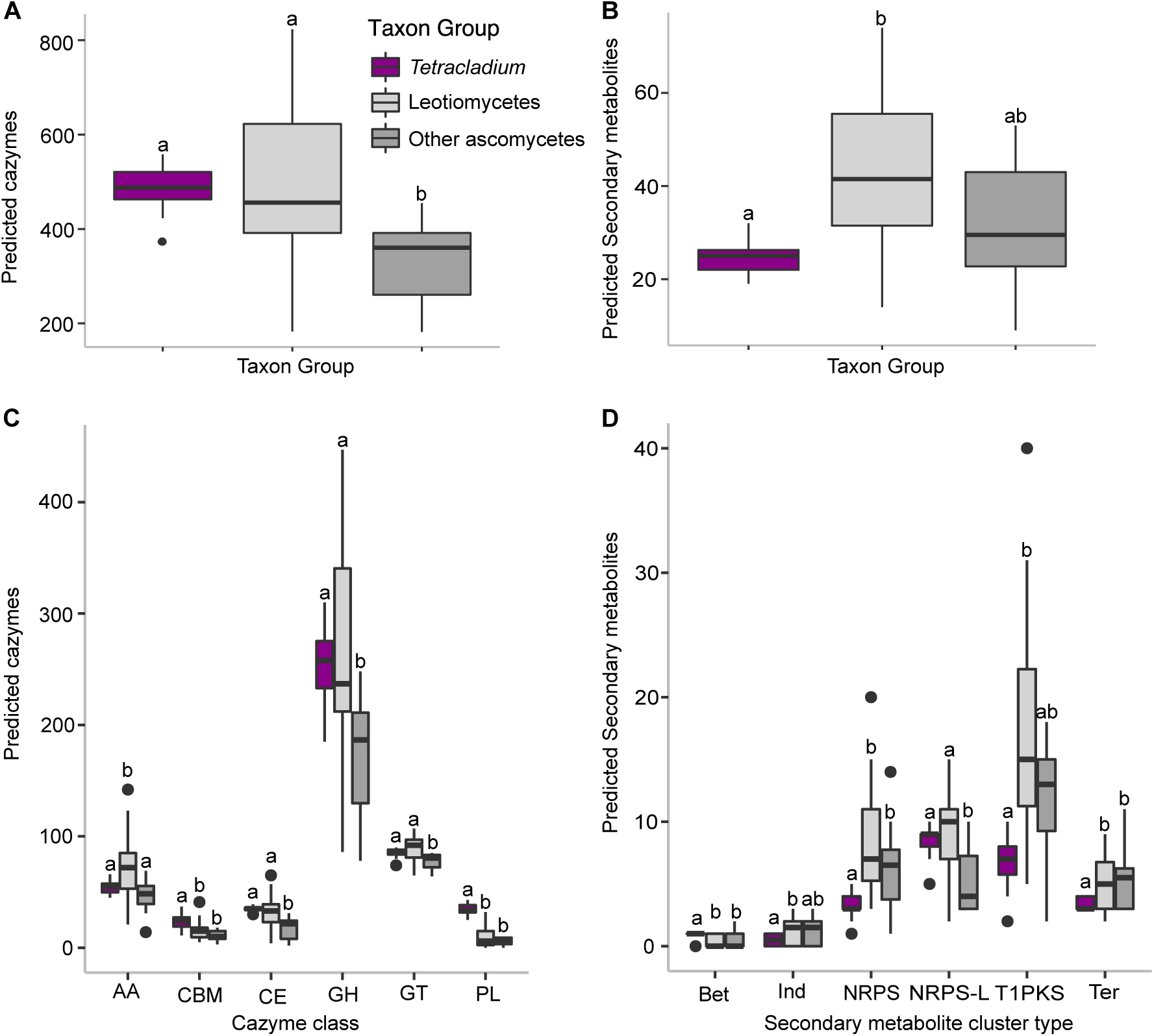
Variation in total number of CAZyme and CBM domains (A), and SM clusters (B) identified in taxon groups containing *Tetracladium* and *Tetracladium*-like genomes (N=24), and the genomes of the other Leotiomycete (N=18) or other ascomycetes (N=8) in the study. Results were analyzed using analysis of variance with Tukey’s HSD for comparisons among taxon-groups. In C and D, separate analyses were performed for each CAZyme or SM type. Taxon-groups sharing the same letter are not statistically different (*p* > 0.05). CAZyme classes and associated modules: Auxiliary Activities (AA), carbohydrate binding modules (CBM), carbohydrate esterase (CE), glycoside hydrolase (GH), glycosyltransferase (GT), and polysaccharide lyase (PL). SM cluster types: betalactone (Bet), indole (Ind), nonribosomal peptides (NRPS), NRPS-Like (NRPS-L) and terpene (ter). Other SM categories did not differ by taxon group (see Table S10).

*Tetracladium* genomes are specifically enriched for polysaccharide lyase (PL) domains relative to other Leotiomycetes and ascomycetes (Fig. 5C; PL: F_2_ = 81.78, *p* < 0.0001; Tukey’s HSD, p ≤ 0.05). *Tetracladium* genomes have 34 ± 4.98 (mean ± 1 standard deviation) PL domains per genome, which is about 3 times the number in other Leotiomycetes (9.95 ± 9.97), and 6 times as many as the other ascomycetes (5.5 ± 3.8). This difference in part reflects the higher copy number of pectate lyase (77) PL3-2 in the *Tetracladium* genomes (9.3 ± 1.4) than the Leotiomycetes (1.8 ± 2.2) and other ascomycetes (0.9 ± 0.8; F_2_ = 134.8, *p* < 2e-16; Tukey’s HSD, *p* ≤ 0.05; Table S6).

*Tetracladium* genomes also contain more CBMs (24.1 ± 6.2), than other Leotiomycetes (15.9 ± 9.1) and ascomycetes (10.6 ± 5.2; CBM: F_2_ = 12.89, *p* < 0.0001; Tukey’s HSD, p ≤ 0.05). Most CBMs (75-91%) were associated with glycoside-hydrolases (GH). Up to three CBMs were found to co-occur flanked by GH32 (GH32+CBM38+CBM38+CBM38+GH32). *Tetracladium* genomes contain more copies of GH32 (4.3 ± 1.4) and CBM38 (2.5 ± 1.6) than the other taxon-groups (Leotiomycetes: GH32 2.6 ± 1.7, CBM38 0.8 ± 1.1; ascomycetes: GH32 2.3 ± 1.8, CBM38 0.3 ± 0.5; F_2_ = 8.4, *p* = 0.0007; Tukey’s HSD, *p* ≤ 0.05). GH32 family enzymes can function as invertases that convert sucrose into fructose and glucose and act on inulin and fructose (77) and CBM38 has inulin-binding function (77). The GH32+CBM38+CBM38+CBM38+GH32 conformation is unique within the genus *Tetracladium* in this study. Versions of this CAZyme with one and two CBM38 between the flanking GH32s were also predicted for some Leotiomycetes (Table S8). All *Tetracladium* genomes except *T. ellipsoideum* and *T. sp*. T11Df had at least one GH32+CMB38 predicted CAZyme.

Copy number of CBM24 also contributes to the difference in CBM among taxon groups; there are 1-15 copies (7.6 ± 4.7) in each *Tetracladium* genome, while the Leotiomycetes (0-11, 3.4 ± 4.0) and ascomycete genomes (0-5, 2 ± 1.8) contain fewer (Table S6). CBM24 has α-1,3-glucan/mutan binding function (77). In *Tetracladium*, CBM24 was almost always predicted in 1-3 copies in association with GH71 an α-1,3-glucanase (77). GH71 is enriched in *Tetracladium* genomes (7.7 ± 3.1) relative to the other Letiomycetes (4 ± 2.4) and ascomycetes (3.9 ± 2.5; F_2_ = 10.39, *p* = 0.0002, Tukey’s HSD, *p* ≤ 0.05). The high copy number of GH71, and its association with 1-3 copies of CBM24 per predicted CAZyme, explains the abundance of CBM24 in the *Tetracladium* genomes.

Strains of the same species have similar predicted CAZyme repertoires overall (Table S9). The same copy number was predicted for 63-93% of the predicted CAZyme domains in the genomes of the the six species represented by more than one strain. This includes *T. sp*. 82A210 within *T. furcatum*, and *T*.*sp*. AFCN889 and *T. sp*. AFCN900 as one species. *Tetracladium furcatum* had the lowest similarity among strains for copy number, which is consistent with the phylogenetics results; F11883 and 82A210 are more similar to each other than to F06983. F11883 and 82A210 are 91% identical for copy number. Chromosome level assemblies are required to determine absolute numbers present in the genomes. However, consistent counts between related genomes can aid interpretation overall and give a first estimate of variation within species. In most cases where counts differed between strains within species the count differed by ±1 (75%, Table S9). Larger differences in copy numbers can be seen for specific CAZymes within species in some cases. For example, GH71 (above) is predicted in 4 copies in *T. marchalianum* F26399 and in 13 copies in *T. marchalianum* JA005 (Table S6). As expected, CBM24 copy number also differed between these two strains (3 and 10 respectively).

### Secondary Metabolites

*Tetracladium* genomes contain 24.7 ± 3.6 SM clusters (Fig. 5B) detected by antiSMASH which is fewer than other Leotiomycetes (43 ± 17.5), but in line with other ascomycetes (31.8 ± 15; F_2_ = 11.53, *p* < 0.0001; Tukey’s HSD *p* < 0.05). All SM data are available in Table S9. *Tetracladium* genomes each contain one betalactone cluster (1.0 ± 0.2), except *T. sp*. T11Df with none, setting them apart from the other taxon-groups (Fig. 5D; Leotiomycetes 0.4 ± 0.5; ascomycetes 0.5 ± 0.8; F_2_ = 9.3, *p* < 0.001; Tukey’s HSD *p* < 0.05). Indole clusters are predicted in only 50% of *Tetracladium* genomes and never more than 1 per genome (0.5 ± 0.5), which is fewer than in Leotiomycete genomes (1.3 ± 1.1) but not different from other ascomycetes (1.3 ± 1.2; F_2_ = 5.1, *p* < 0.01; Tukey’s HSD *p* < 0.05). More nonribosomal peptides (NRPS) are found in Leotiomycete (8.2 ± 4.6) and other ascomycete (6.5 ± 4.1) genomes than in *Tetracladium* genomes (3.2 ± 0.8; F_2_ = 12.7, *p* < 0.0001; Tukey’s HSD *p* < 0.05). All Leotiomycete genomes (9.2 ± 3.7), including *Tetracladium* (8.4 ± 1.2), contain more NRPS-like SM clusters than the other ascomycetes (5.3 ± 2.8; F_2_ = 6.4, *p* = 0.004; Tukey’s HSD *p* < 0.05). Fewer Type 1 polyketide synthase (T1PKS) are predicted in *Tetracladium* genomes (6.8 ± 2.0) than in leotiomycetes (17.2 ± 9.1), but neither group differs from the ascomycetes (11.6 ± 5.1; F_2_ = 15.66, *p* < 0.0001; Tukey’s HSD *p* < 0.05). Both Leotiomycete (5.1± 2.0) and other ascomycete genomes (5.5 ± 2.7) contain more terpene SM clusters than *Tetracladium* genomes (3.4 ± 0.5; F_2_ = 7.7, *p* = 0.001; Tukey’s HSD *p* < 0.05). The frequencies of ribosomally synthesized and posttranslationally modified peptides (fungal-RIPPs), phosphonates, siderophores and Type III polyketide synthases (T3PKS) clusters were low and did not differ among taxon groups (Table S10).

While *Tetracladium* genomes each contain around 24 SM clusters detectable by antiSMASH only 13% returned a BLAST match for most similar known cluster in the MiBIG database(78). In total 18 SM clusters were identified to type (Fig. S4, Table S10). A nonribosomal peptide synthetase (NPS), Dimethylcoprogen, was the only SM predicted in all *Tetracladium* genomes (100% similarity (75)). This cluster is found in only 2 of 18 (11%) of the other Leotiomycete genomes and 3 of 8 (38%) other ascomycetes. All other identified SM clusters were Group or taxon specific within *Tetracladium* (Table S10). Unique to some members of Group A: Solanapyrone, Hexadehydro-astechrome, Cytochalasin, Phyllostictine A/phyllostictine B, Citreoviridin, Clavaric acid, Aureobasidin A1 and Chaetoglobosins. Unique to Group some members of Group B: Clapurines, Naphthopyrone, Shearinine D, PR toxin, Pyranonigrin E, Azanigerone. Unique to Group C: Duclauxin. Depudecin (Groups A and B), Brefeldin (B and C) were also predicted.

## DISCUSSION

### Relationships among *Tetracladium* and *Tetracladium*-like fungi

This study presents the first phylogeny with all species of *Tetracladium* described to date. The eleven described species and all newly sequenced *Tetracladium*-like (putative) strains/species form a monophyletic group with three partitions (Fig 2, Groups A, B, and C). Whereas interspecific relationships within *Tetracladium* were unresolved in analyses using 18S (79), 28S (20), and ITS+28S (80) data and few taxa, relationships within the genus are largely stable across analyses and datasets herein. Within the *Tetracladium* and *Tetracladium*-like group only the branching of *T. furcatum* and *T. maxilliforme* varied between analyses. *Tetracladium furcatum* and *T. maxilliforme* are well supported separate species, but it is unclear whether they are sister species or whether *T. furcatum* alone is sister to the rest of Group A.

The majority of species and strains in Group A originate from submerged plant debris or foam that forms on rivers and produce stereotypical *Tetracladium*-shaped conidia which are typically distinctive between species (19, 38). Although conidia have not been observed for the strain isolated as an endophyte of wheat (Sweden, 2007), *T. sp*. 82A210, this strain falls within the species *T. furcatum*, and is highly similar to strain F11883 (Czech Republic, 1983) which was isolated from foam as a typical aquatic hyphomycete (Table S1). This result supports the idea that *T. furcatum* is multifunctional, as opposed to having different ecologies for morphologically similar species. *Tetracladium furcatum* was first reported as a root endophyte in 1996, based on the morphological identification of conidia (81). It has also been associated with the endophytic lifestyle in metabarcoding studies, including studies of terrestrial plants from the high Arctic (35) and submerged aquatic plants in Norway (15). *Tetracladium nainitalense* which was isolated as endophyte of *Eupatorium adenophorum* (11), is most closely related to species isolated as aquatic hyphomycetes and has itself been isolated from foam in a stream (morphological identification (82)). It should be noted that *Tetracladium* species are found as endophytes even in non-stream, non-riparian habitats. This study also confirms the phylogenetic position of T. sp. F10008, collected as an aquatic hyphomycete from Malaysia in 2008, as the sister species of *T. apiense* (19).

In the majority of cases when more than one strain per species was sequenced the strains are resolved together as expected. Most strains in Group A were isolated by experts in aquatic hyphomycetes (Table S1), who will have relied on spore morphology for initial identification. This suggests that morphological identification of most Group A species from field samples can be reliable. However, two species are potentially problematic. Strains historically identified as *T. marchalianum*, F12812 (called *T. sp*. F12812 herein) and F26399, are not conspecific. The same is true of two strains of *T. breve* (F12505 and F10501). Rather, the *T. marchalianum*-like strain F12812 is the sister species of the *T. breve*-like F12505, and *T. marchalianum*-like F26399 is sister to *T. breve*-like F10501, and the two pairs are divergent. This finding is consistent with previous studies based on one or few genes (19, 83). The *T. marchalianum* and *T. breve*-like strains also differ in predicted secondary metabolite profiles (Table S10), suggesting that secondary metabolite profiles might be valuable tools for species identification in the same way that protein fingerprinting is being developed (51). *Tetracladium marchalianum*-like strains can readily be catagorizied as F12812 or JA005/F26399-like using beta-tubulin sequence clustering (SI Fig. X), based on a preliminary analysis of strains identified as *T. marchalianum* from a population genetics study (84). Note, the strains from that study all cluster with JA005/F26399(Fig. S5). Both *T. marchalianum* and *T. breve* require further study and taxonomic revision. Taxonomic revision is beyond the scope of the work presented here, but is ongoing

As more fungi related to Groups B and C are discovered it is probable that these groups will be described as separate genera. In addition to being divergent from Group A in the phylogeny, the fungi in Group B have not been observed to produce the conidia typical of the genus. *Tetracladium elipsoideum* and *T. globosum* are named for their ellipsoid and globose conidia. *T. psychrophilum*, although described without conidia, does produce floating multiseptate elongated allantoid-lunate conidia when submerged (Anderson personal observation). Sporulation in the two endobryophytic strains (*T. sp*. AFCN889 and AFCN900) has not been observed. Based on strains included here, including the ITS phylogeny below, it appears that Group B may predominantly contain psychrophilic or psychrotolerant fungi. The sole representative of Group C, *T. sp*. T11Df, is particularly unusual in that it grows as a yeast, producing short pseudohyphae in culture. All other species in the genus are known in filamentous forms. Spores have not yet been observed in this species.

The family level phylogenetic relationships resolved in this study are in line with the results from previous studies (85-87). Within the Leotiomycetes, all clades identified by Johnston and colleagues (88) and were represented here, are recovered. The family containing *Tetracladium*, Vandijckellaceae (88, 89), was included in an analysis based on 15 concatenated sequences (88). The tree herein differs from that result in the branching order of the Vandijckellaceae and helotioid taxa. However, the corresponding nodes are not well supported in the Johnston tree where they receive maximum 93% UFBoot, but are fully supported herein (100% UFBoot). The minimum support considered reliable in ultrafast bootstrapping (UFBoot) in IQ-TREE is 95% (90, 91).

### Broad ecological and geographical diversity in *Tetracladium*

*Tetracladium*-like fungi are increasingly being reported from aquatic and terrestrial sources, including from soils and as endobionts of plants, from around the world. Identification as “*Tetracladium*-like” is frequently based only on ITS sequence data and ITS data can be useful to identify species of *Tetracladium* in at least some cases (19). Here, from analysis of the *Tetracladium*-like ITS sequences available in GenBank, it is clear that both ecological and geographical diversity are wide-spread within the genus (Fig. 3). These sequences represent diversity in *Tetracladium* from 32 countries, regions, and territories, from sea level to high elevation, and from the Arctic to the Antarctic (Table S5). It is striking how many *Tetracladium* sequences are coming from polar and alpine regions, high elevations, or are glacier associated (Fig. 3). Frequently, fungi from these extreme environments that were sequenced from soil, water or non-orchid plants are similar in ITS to those from orchids from more temperate climates. Overall, patterns of strains from different sources or lifestyles being distinct from each other are not observed. Rather, in most cases, the ecologies and lifestyles are mixed among the groups of sequences most similar based on ITS. These results suggest that multifuncationality is widespread across the genus. Further, *Tetracladium* fungi are found in association with a broad diversity of plants (Table S5).

### Ecological clues from CAZymes and secondary metabolites

The CAZyme and secondary metabolite profiles of *Tetracladium* species are more similar to each other than to other taxa, regardless of ecology (Fig. 4). This suggests a strong role for phylogeny in shaping the CAZyme and SM content of genomes within the genus. In comparison to genomes of other Leotiomycetes and more distantly related ascomycetes, *Tetracladium* genomes are enriched for PL domains which are associated with degradation of pectin and pectate. *Tetracladium* genomes contain around 3-6 times more PL domains than the other fungi in the study which may be related to their ecological role as aquatic hyphomycetes. Aquatic hypyomycetes are major decomposers of leaves in streams and pectin and pectate are complex polysaccharides that are abundant in leaves. In contrast, the genome of *Xylona heveae*, a horizontally transmitted endophyte of sapwood in rubber trees, contains no predicted PL (92).

*Tetracladium* genomes contain a particularly large number of pectate lyase PL3-2 (EC 4.2.2.2) genes, which is a feature they share with *A. tetracladia*, the only non-*Tetracladium* aquatic hyphomycete in the study (Table S6). The PL repertories of *Tetracladium* species and *A. tetracladia* are very similar overall, however they are also qualitatively similar to the two *Cadophora* species in the study (Table S6). The sequenced strain of *C. malorum*, was isolated from a deep sea shrimp from a depth of 2300 m below sea level at a hydrothermal vent along the Mid Atlantic Ridge (93), the other *Cadophora sp*. was isolated as an endophyte of *Salix rosmarinifolia*; neither is expected to decompose leaves. Thus, determining relative importance of ecology and phylogeny in shaping the PL content of these genomes requires genomes from additional taxonomically and ecologically diverse fungi.

Fungal saprotrophs (of plants), endophytes, and plant pathogens are typically able to degrade plant cell walls in order to enter a plant host or grow through a plant substrate and to obtain nutrition from plants. Thus, some overlap in the CAZyme and SM content in the genomes of fungi that exhibit these ecologies should be expected especially among taxa that are multifunctional (94). PL3-2, the most abundant PL in *Tetracladium* genomes (above), is best known as part of the molecular arsenal of plant pathogens including *Botrytis cinerea* and is highly expressed in developing infections (*e*.*g*. tomato (95)). Also, pectic enzymes can elicit defense responses in plants (96). In *Tetracladium*, expression of PL3-2, and other PLs, may require tight context dependent control, with high expression during saprotrophy to degrade leaves, but no-to-low expression when living as endobionts to avoid triggering plant defenses. The Dimethylcoprogens, nonribosomal peptide synthetases involved in synthesis of siderophores (97) which have iron uptake and storage functions, have also been reported as common among taxa within the Pleosporales, Dothideomycetes (98) an order that includes many plant pathogens. Dimethylcoprogen, and siderophores generally, have been associated with pathogenicity for some fungi including the corn pathogen *Cochliobolus heterostrophus* and wheat pathogen *Fusarium graminearum* (99). However, iron homeostasis and storage have other important roles in fungi, including resistance to reactive oxygen species (97, 99) and the maintenance of mutualism in endophytic fungus–plant interactions (100). Going forward, comparisons expression patterns associated with these PLs and SM clusters can help dissect how organisms with similar molecular “toolboxes” selectively wield these tools appropriately for saprotrophs, endobionts, and pathogens.

*Tetracladium* genomes are also rich in GH71 domains which are α-1,3-glucanases and their associated non-catalytic CBM24 α-1,3-glucan/mutan binding modules, suggesting that these fungi are well-suited to breaking down fungal cell walls; is an important component of the cell walls of filamentous fungi and dimorphic yeast. As was the case for *S. sclerotiorum* and *B. cinerea* when enrichment for GH71 in these fungi was first reported, it is unknown whether the abundant α-1,3-glucanases in these genomes is associated degradation of the fungi’s own cell walls or those of antagonistic fungi(101), or whether these enzymes play other roles entirely. Interestingly, α-1,3-glucan is important in the matrix of fungal and bacterial biofilms (102) which can be disrupted by glucanases (103, 104). Decomposing leaves in streams and the roots of plants are covered by microbial biofilms and it is possible that enrichment for *α*-1,3-glucanase in *Tetracladium* genomes is associated with interactions with those biofilms. Further, there is some evidence that α-1,3-glucanases prevent plant detection of *β*-1,3*-*glucan in in invading pathogens, negatively impacting plant defensive responses. Transgenic rice plants expressing bacterial (105) or fungal (106) α-1,3-glucanases demonstrate protection against fungal pathogens. Thus, it is also possible that these enzymes are important in plant-fungal mutualism and beneficial to plants (107, 108).

*Tetracladium* genomes also contain more GH32 catalytic domains and associated CBM38 modules than other taxon groups studied here. GH32 enzymes can function as invertases and also act on inulin and fructose (77). Given the inulin-binding function of CBM28 the inulinase function may be most enriched in *Tetracladium*. Inulin is a reserve carbohydrate in some plants that is stored in roots, taproots, and bulbs. Some strains/species of *Tetracladium* were found to have these components in a GH32+CBM38+CBM38+CBM38+GH32 conformation; unique among the taxa in this study. These observations suggest that *Tetracladium* species obtain nutrition from inulin when living as endobionts of plants. Studies of *Tetracladium* species as root endophytes are needed to test this hypothesis.

## CONCLUSIONS

The genomes of 24 *Tetracladium* and *Tetracladium*-like fungi, including representatives of all described species, were sequenced and used to resolve relationships among the taxa and to improve our understanding of ecological and genomic diversity in this group of ecologically important, multifunctional fungi. All genome-sequenced *Tetracladium* and *Tetracladium*-like fungi in this study form a monophyletic group, which may in time be subdivided into separate genera. From analysis of ITS sequences from water, soil, and plants from around the world, it emerges that multifunctionality may be widespread throughout the genus, that many species have multifunctional lifestyles. Further, *Tetracladium* is frequently sampled from extreme and cold environments, suggesting that these fungi may have important roles in those ecosystems and also may produce secondary metabolites or enzymes of interest for industrial applications. Studies are needed to investigate the terrestrial and endiobiont roles of *Tetracladium* fungi, including of where in plant roots these fungi are found, whether they utilize inulin as a source of nutrition as endobionts, and how PL expression is controlled in saprotroph and endiobiont contexts. Lastly, these fungi are more similar to each other in genome content for SMs and CAZymes than they are to other taxa, regardless of variation in the ecology of the fungi, suggesting that within *Tetracladium*, broad ecological diversity and multifunctionality can be achieved among taxa using highly similar genomic toolkits.

## ACKNOWLEDGEMENTS

We thank Iker Irisarri and Dan Vanderpool for advice on phylogenomics and annotations. We thank Anna Rosling for reagents and Doug Scofield, Ioana Onut Brännström, and Diem Nguyen for analysis support. We are grateful to Monika Laichmanová and the CCM, Marcelo Baeza, Dai Hirose, and Magdalena Grudzinska-Sterno for cultures. Sequencing was performed by the SNP&SEQ Technology Platform in Uppsala. This research, and J. Anderson, are funded by grant #2016-03595 from Vetenskapsrådet, The Swedish Research Council.

---

**Figure S1.**
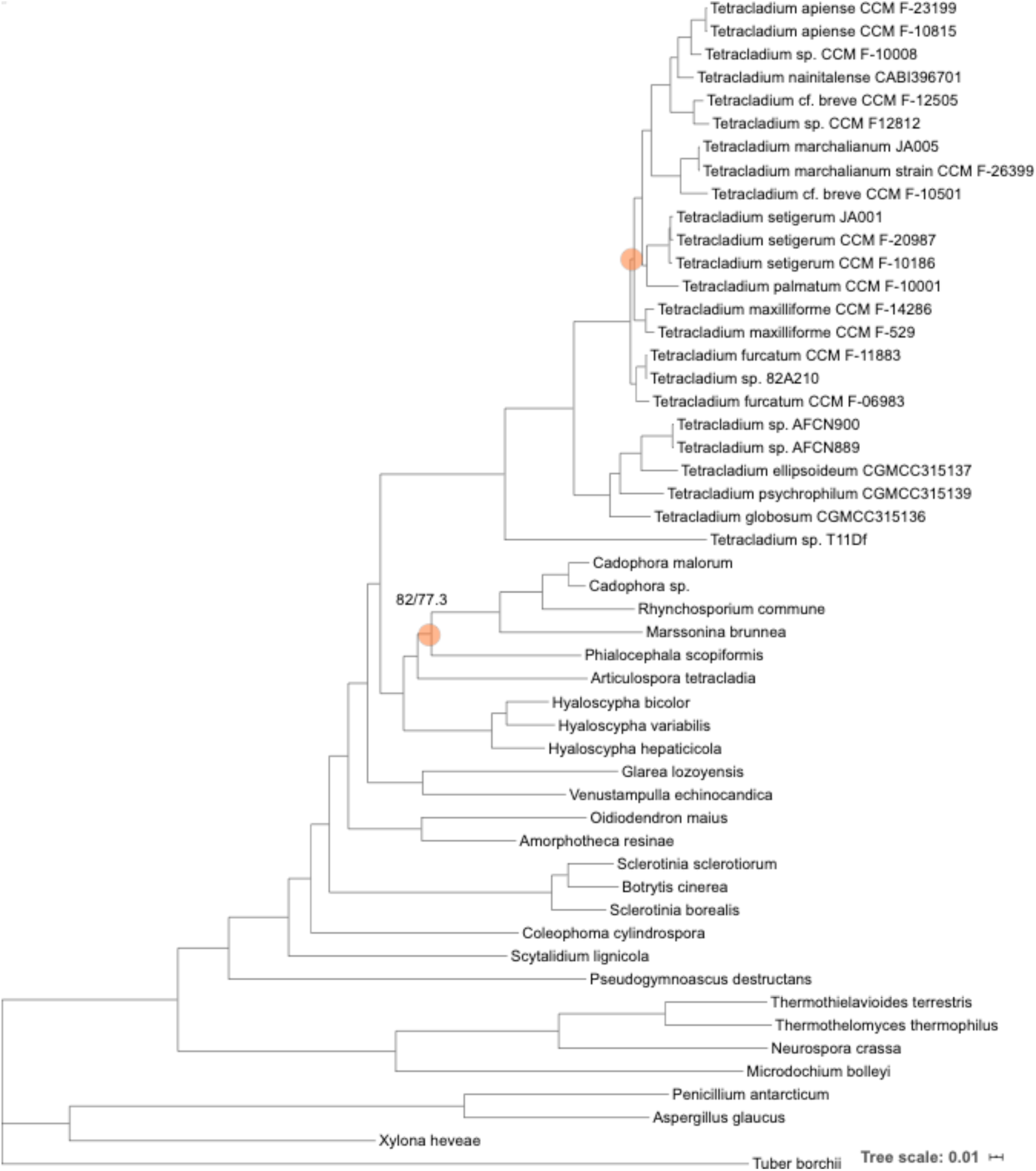
ML tree from analysis of the all-SCO dataset with model LG+F+I+G4. All branches received UFBoot values > 95% and SH-aLRT > 80% unless indicated (UFBoot%/ SH-aLRT%). The tree scale represents the expected number of nucleotide substitutions per site.

**Figure S2.**
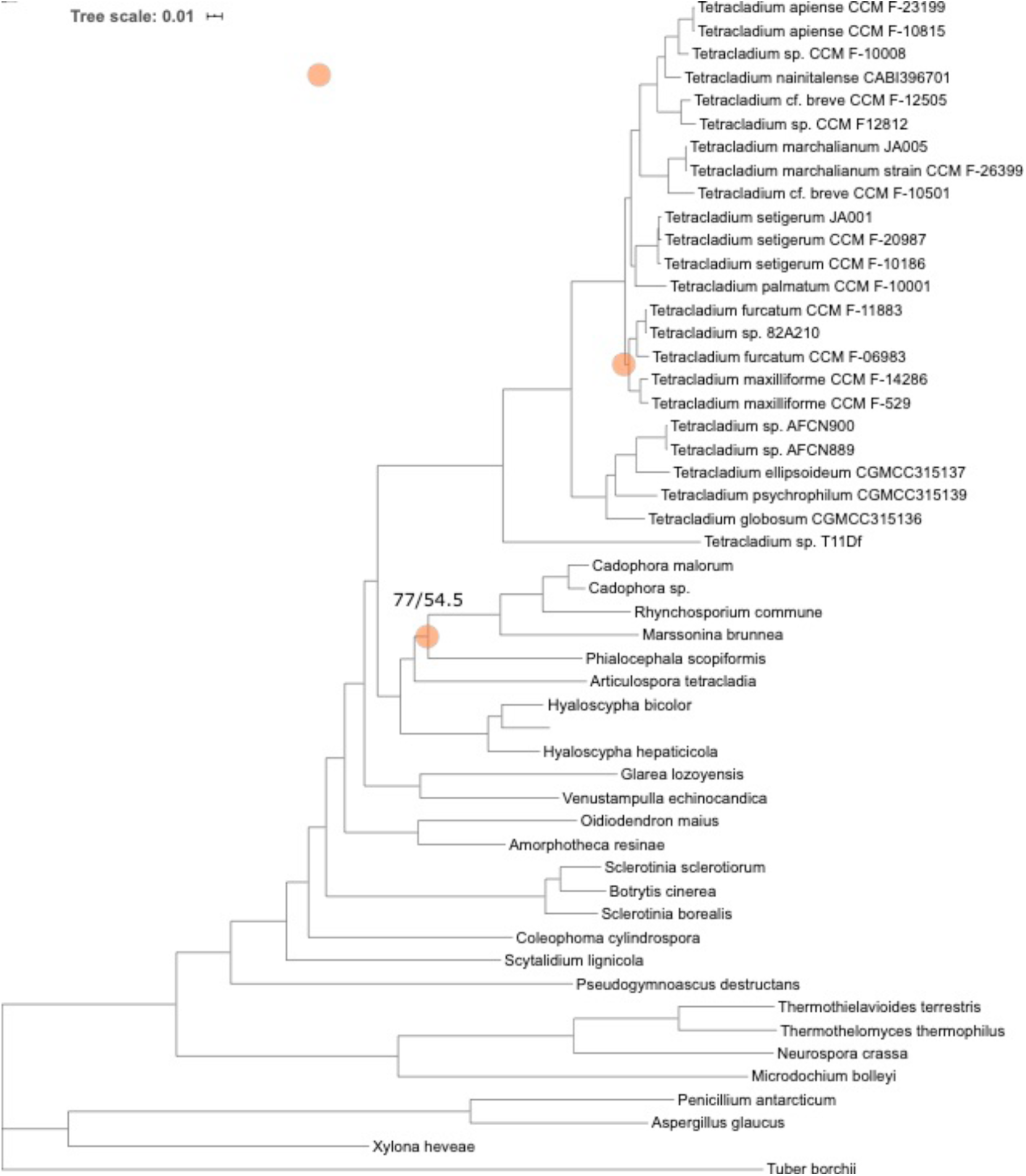
ML tree from analysis of the asco-SCO dataset with model LG+F+R6. All branches received UFBoot values > 95% and SH-aLRT > 80% unless indicated (UFBoot%/ SH-aLRT%). The tree scale represents the expected number of nucleotide substitutions per site.

**Figure S3.**
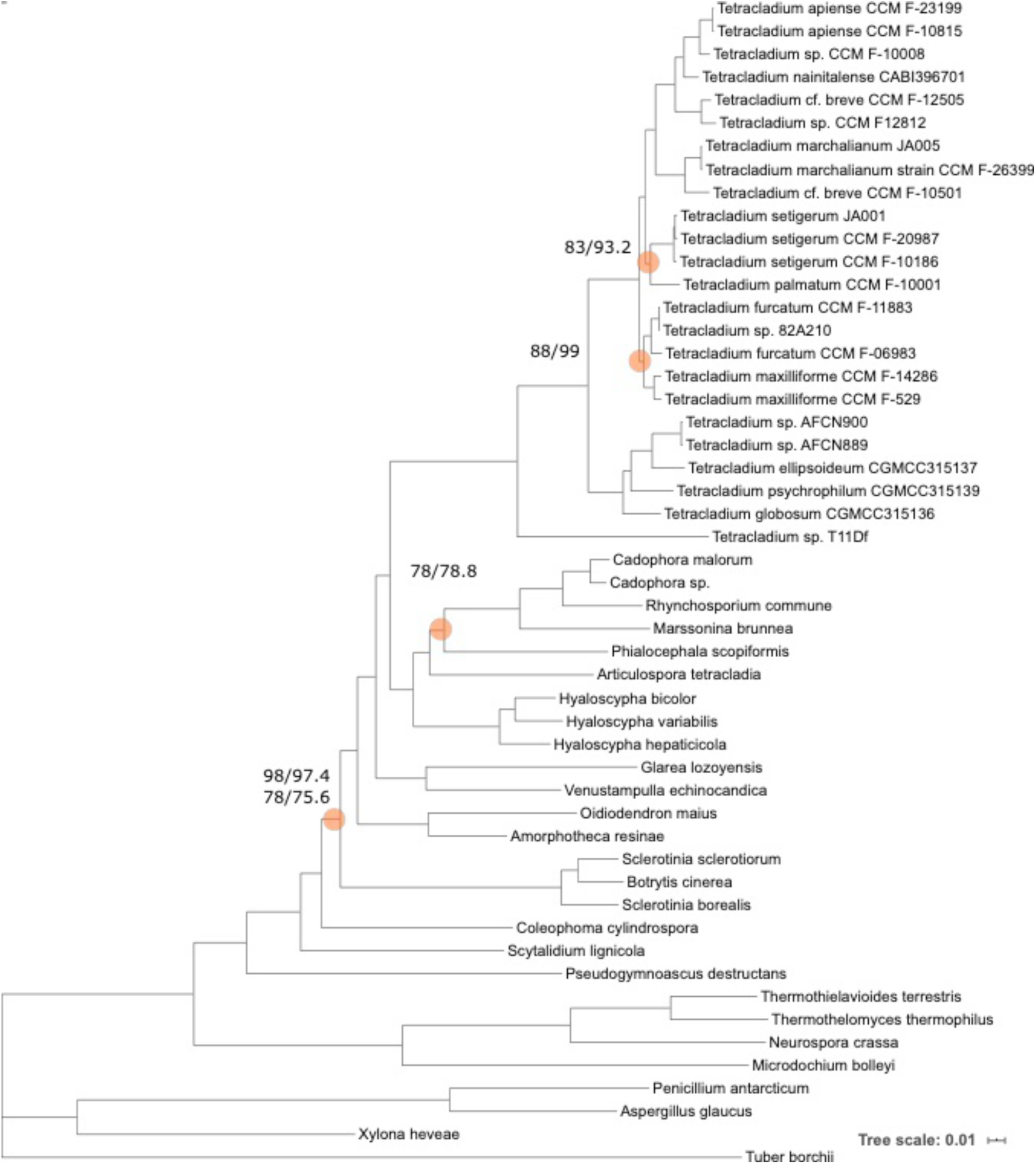
Summary of ML trees from analysis of the fungi-SCO dataset model LG+F+I+G4 and using the FreeRate model. All branches received UFBoot values > 95% and SH-aLRT > 80% unless indicated. Support values are presented as UFBoot%/ SH-aLRT% for LG+F+I+G4 above, the FreeRate model below. Tree scale refers to the FreeRate model analysis. The tree scale represents the expected number of nucleotide substitutions per site.

**Figure S4.**
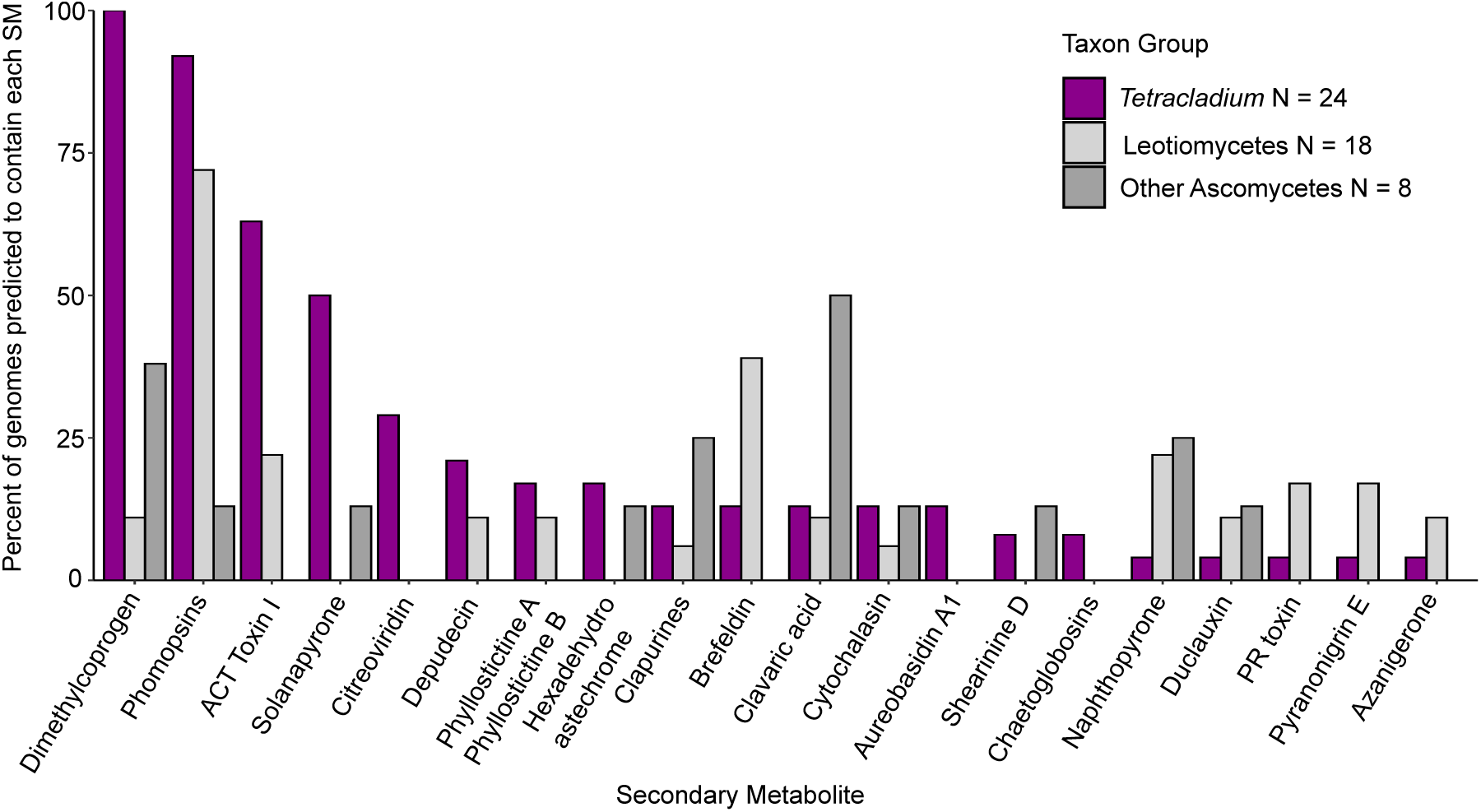
The 18 secondary metabolite clusters in *Tetracladium* genomes that were identifiable to type and the percent of genomes in each taxon-group where each SM type was also identified. Note the difference in sample size among taxon-groups.

**Figure S5.**
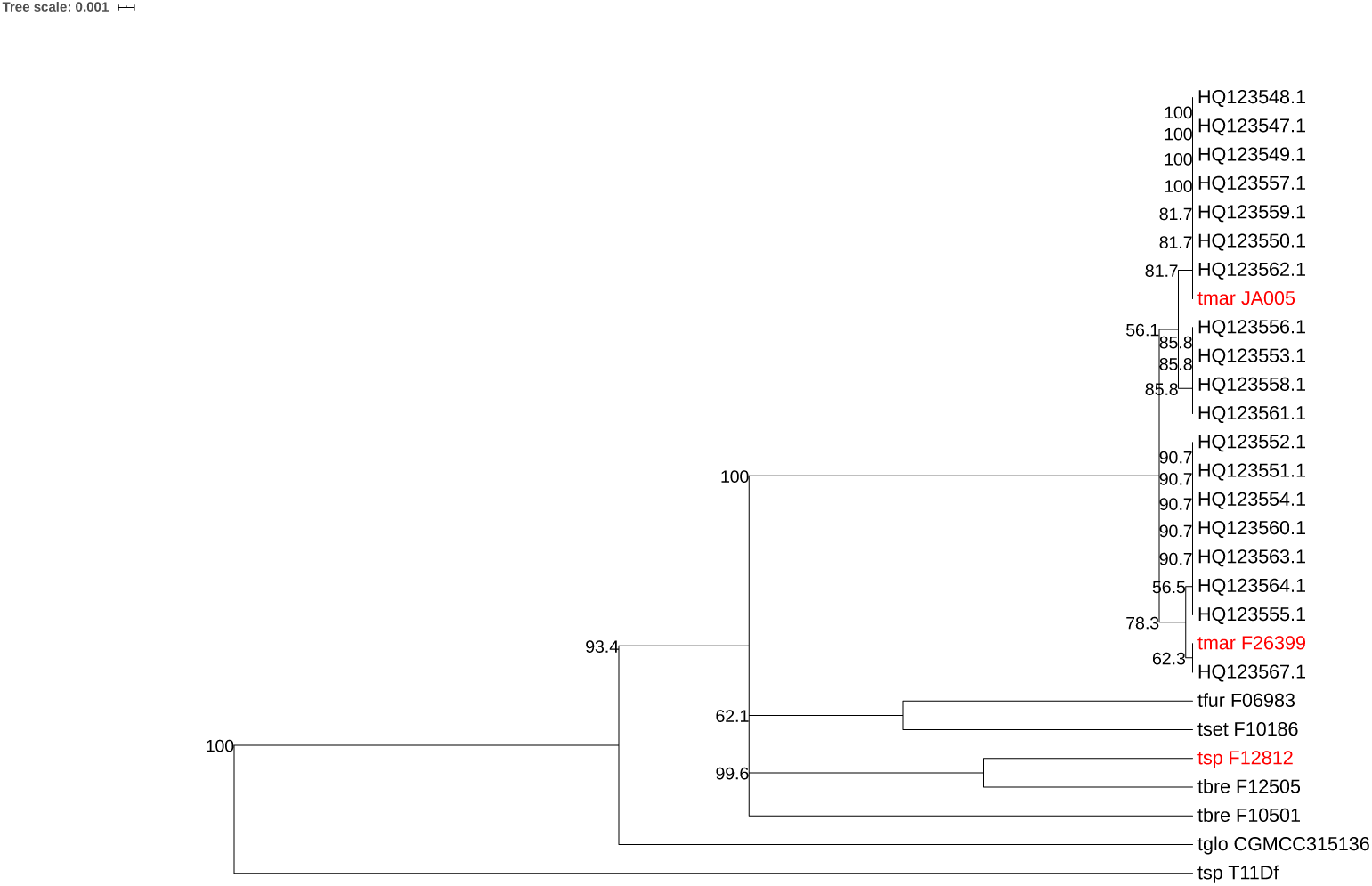
UPGMA clustering of 46 beta-tubulin sequences. Analysis performed in Geneious 10.2.4 using a Jukes-Cantor genetic distance model branch support estimated using 1000 bootstraps with resampling. All sequences with GenBank accessions starting with HQ come from the population genetics study of Anderson and Shearer (84).

